# The TRX assay for triplet repeat expansions

**DOI:** 10.1101/2024.12.18.629104

**Authors:** Thomas O’Brien, Karolina Salciute, Shaylee Kieffer, Nehaa Kandasamy Packil Ponraj, Madeleine Richer, Paulina Marzec, Caroline Benn, Darren G. Monckton, Vincent Dion, Robert S. Lahue

## Abstract

Expansion mutations of triplet repeat sequences cause numerous inherited neurological diseases. In some diseases, affected individuals display somatic expansions in affected tissues which have been linked to accelerated disease onset and progression. There is currently considerable interest in developing therapies to slow somatic repeat expansions to delay or block disease onset. *In vitro* assays are particularly important to evaluate potential therapeutic interventions. Current assays typically use physical methods to monitor triplet repeat lengths within a population of cells. While useful, most of these assays are relatively slow (∼six weeks) and are somewhat limited in sensitivity to rare events. Here, a new assay, called TRX, is described to monitor CAG•CTG triplet repeat expansions more rapidly and with better sensitivity. TRX uses human tissue culture cells expressing two fluorescent proteins. Red fluorescent protein TagRFP658 is constitutively expressed and serves as an internal control. GFP is expressed in a CAG•CTG repeat length-dependent manner, with longer repeat lengths predicted to give higher green fluorescence intensity. Standard flow cytometry allows quantification of changes in fluorescent signal as a simple readout with <2% sensitivity. Two independently derived cell lines with 63 or 59 CAG repeats yielded similar rates of TRX activity. Cells with increased green fluorescence were observed within one to two weeks of culture, with longer times leading to additional signal. The appearance of green fluorescence was partly dependent on MutSβ, the DNA MSH2-MSH3 complex, based on siRNA knockdown of MSH3. However, physical analysis of the CAG•CTG repeat tracts by MiSeq deep sequencing or capillary electrophoresis showed limited changes in the length of the repeat tracts. We conclude that the TRX assay is a promising new tool for monitoring CAG•CTG repeat expansions but that further development of the assay is needed to make it fully useful.

## Introduction

DNA triplet repeats are three-base tandem arrays that can mutate by either expansion (gain of repeats) or contraction (loss of repeats). Expansion mutations are of medical interest because inherited expansions cause at least 20 debilitating neurological diseases [1,2]. For some of these diseases, the repeat tract continues to expand somatically in affected tissues [3,4]. Recent evidence has shown that these somatic expansions contribute significantly to the age of disease onset [5–7] and to the rate of disease progression for some triplet repeat disorders [4]. This development has spurred interest in potential therapies aimed at inhibiting somatic expansions to delay or block disease onset [8]. This therapeutic strategy is enhanced by the fact that key target proteins driving somatic expansions have already been identified through human genetics and model system studies [4,9–12]. These proteins include the mismatch repair factors MSH3, MLH1, PMS1, and MLH3, whose ablation or inhibition reduces expansion rates and delays disease in model systems [13–15]. Thus, the targets for therapy are already identified. A second key fact is that expansions of the triplet repeat sequence CAG•CTG underlie at least 15 of these inherited disorders, and several diseases in this category display somatic expansions [16]. If therapies can be found that slow somatic expansion and are effective for one disease, those same therapies might be applicable to other CAG•CTG expansion disorders. These findings help justify a concerted effort currently aimed at finding therapeutic approaches that target somatic triplet repeat expansions.

The discovery process for therapies relies on *in vitro* assays as an important step in drug development. A common method currently used to measure somatic expansions couples PCR amplification across the repeat tract with capillary electrophoresis. For examples, see [17–19]. The resulting “hedgehog” pattern results from the combination of repeat length diversity in the cell population as well as PCR slippage events. The modal peak height is typically used to monitor changes in repeat length in the population. In some cases, a somatic instability index is calculated to include weighting of non-modal peaks [20]. This approach is useful, although it typically requires ∼six weeks of culture time to detect significant changes to repeat tracts. It is also only moderately sensitive since numerous cells in the population must contain expansions to move the pattern detectably. The somatic instability index improves sensitivity somewhat. A second method uses ultradeep MiSeq sequencing to display somatic expansion patterns [21,22]. This method is very sensitive for small length changes, but it is not sensitive to rare large length changes, and its speed is limited by the time required for sequencing sample preparation and analysis. Cell-based fluorescent assays exist where contractions of CAG•CTG tracts cause increases in GFP signal [23,24], but expansions lead to a decreased GFP signal [25]. There is space to develop a new in vitro assay for triplet repeat expansions with the goals of improving speed, sensitivity, and throughput. This manuscript describes the TRX assay as a new assay method aimed at achieving these improvements.

## Materials and methods

### Cell culture

HT-1080 cells were obtained from the Leibniz Institute DSMZ (#ACC 315; [26]). Cell identity was confirmed by simple tandem repeat analysis (**Supplemental Fig. 1A**). The cells were also tested by western blotting and showed positive signals for MSH2, MSH3 and MSH6 (**Fig. 3B**), as well as MLH1, PMS2, and HPRT (data not shown). HT-1080 and all its derivatives described here were cultured in DMEM low glucose medium supplemented with 10% fetal bovine serum and 1% Pen Strep (all from Sigma) at 37°C and 5% CO_2_. When present, doxycycline hyclate was added to a final concentration of 1,200 ng/ml.

### Targeting plasmids for insertions into HPRT

Two targeting plasmids were created for inserting cloned sequences into the *HPRT* gene on the X chromosome. Each plasmid is based on pUC19 and carries two ∼600 bp homology arms. In between the arms is a unique NotI site for cloning in fluorescent reporters for the TRX assay. The first vector, pUC19_HA1, is designed to target CRISPR/Cas9-mediated insertions into intron 2 of *HPRT* at coordinates NC_000023.11:g.134473748. The second vector, pUC19_HA2, targets insertions into intron 5 of *HPRT* at coordinates NC_000023.11:g.134490280. The two integration sites are 16,531 bp apart. The two targeting plasmids were created synthetically by a commercial vendor (GenScript) and confirmed by DNA sequencing. See **Supplemental Figure 2** for details.

### GFP vector pTOB.041

Vector rAAV.TetR-KRAB/d2GFP [27] was kindly provided by Dr Caroline Le Guiner Blanvillain (Univ. Nantes, France). This construct contains the elements TetO through polyA site (except for the CAG tract and *Pem1* intron) shown in **Fig. 1B**. Vector rAAV.TetR-KRAB/d2GFP was cut with restriction enzymes XhoI, SfiI, and ClaI. The DNA ends were converted to blunt ends by treatment with T4 DNA polymerase and dNTPs. DNA fragments were separated by preparative agarose electrophoresis and a 4.6 kb band was excised and purified. Insertion vector pUC19_HA1, targeting *HPRT* intron 2, was cleaved with NotI, followed by creation of blunt ends, and gel purification as above. Insert and vector DNA were ligated and transformed into *E. coli*. Colonies were screened by PCR using primers 040.05 and 040.06 to detect the *GFP* gene (primer sequences are found in **Supplemental Table 2**). Miniprep DNA from six positive colonies was screened by *Ase*I digestion and three clones gave the predicted bands of 3,455 and 3,001 bp. One clone, named pTOB.025, was selected for further development.

**Figure 1.**
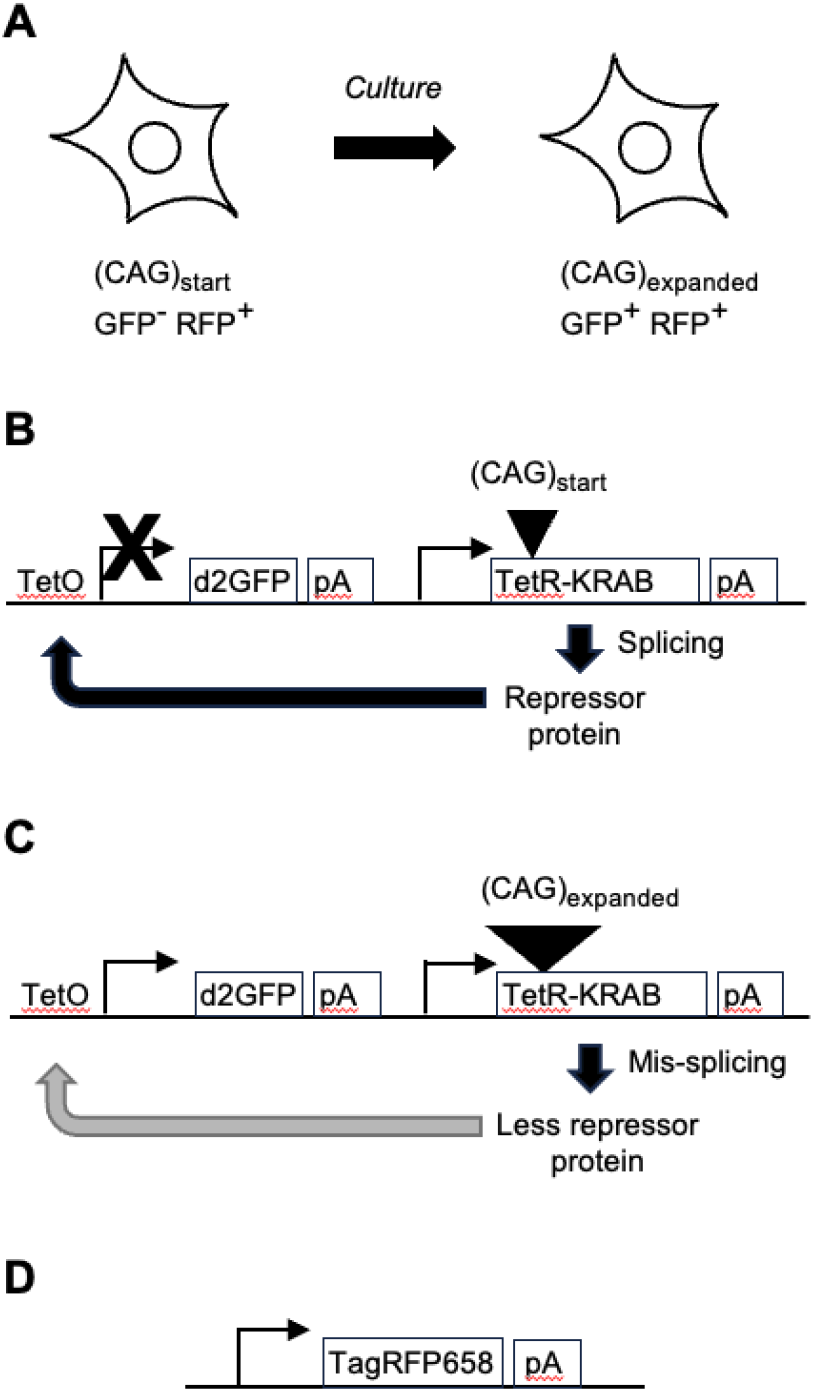
Schematic diagram of TRX assay. **A**, Concept: the short CAG tract in the starting cell does not allow expression of GFP (GFP-), whereas the TagRFP658 tag is expressed constitutively (RFP+). Upon CAG repeat expansion, the fluorescent phenotype switches to GFP+ RFP+. **B**, Starting cell: the gene for the repressor protein TetR-KRAB contains an artificial intron (triangle) containing the starting-length CAG repeat. Splicing of the intron is efficient, allowing production of the TetR-KRAB protein, binding to TetO and silencing of d2GFP (X). The cells are correspondingly GFP-. **C**, Cell with expansion: TetR-KRAB transcripts are predicted to be mis-spliced due to the longer CAG tract, lessening the abundance of TetR-KRAB protein and reducing silencing at TetO (gray arrow), allowing d2GFP expression and a GFP+ phenotype. **D**, All cells constitutively express TagRFP658 and are therefore RFP+. pA, polyadenosine tract.

To insert the *Rattus norvegicus Pem1* intron [24] into pTOB.025, plasmid pVIN110 was PCR amplified with Q5 DNA polymerase and primers oTOB.105 and oTOB.106. The 1,536 bp amplicon was gel purified and mixed with pTOB.025 that had been cleaved *in vitro* with Cas9 and gRNA 2 according to the manufacturer’s specifications (NEB, followed by gel purification of the linear 8.9 kb product. The two DNA molecules were spliced by Gibson assembly according to the manufacturer’s specifications (New England Biolabs), transformed into bacteria, and colonies were screened by PCR using primers pTOB109.02 and pTOB109.03 to look for a 232 bp product internal to *Pem1*. Six positive candidates were examined by HindIII digestion and found to have the predicted digestion pattern. One clone, named pTOB.036, was used going forward.

To insert CAG repeats into the *Pem1* intron of pTOB.036, this plasmid was cleaved *in vitro* by Cas9 plus gRNAs 7 and 8 together. Control reactions used single gRNAs. Preparative gels indicated double-cutting when both gRNAs were present, and the resulting 9,345 bp product was purified. The cleaved plasmid was mixed with a PCR product of the *Pem1* intron containing 123 CAG repeats and assembled using the Gibson protocol. Resultant bacterial colonies were screened by PCR using primers pTOB.109.02 and pTOB.109.03. While some clones gave the expected product size of ∼750 bp, corresponding to the presence of 123 CAG repeats, other colonies showed smaller amplification products, presumably due to spontaneous repeat contraction events in *E. coli*. NotI digests of the candidate plasmids indicated clones with tracts of approximately 50 to 121 CAG repeats. This series of plasmids was designated pTOB.041, followed by the number of CAG repeats as determined by Sanger sequencing. Plasmid pTOB.041.59 was found to have 59 uninterrupted repeats, and plasmid pTOB.041.063 contained a CTG interruption in triplet 15 to yield a sequence of (CAG)_14_CTG(CAG)_48_.

### Creation of pTOB.033 with TagRFP658 into HPRT intron 5 targeting vector

Template plasmid 178974 (Addgene) was amplified by Q5 DNA polymerase (NEB) using PCR primers oTOB.144 and oTOB.145. The resulting product was spliced into NotI-cut pUC19_HA2 by Gibson assembly (NEB). Following transformation into *E. coli*, single colonies were screened by PCR using primers oTOB.029, oTOB.032, oTOB146 and oTOB.148 to detect splice junctions. Six plasmid clones with positive PCR signals were transfected transiently into HT-1080 cells and flow cytometry was used to detect red fluorescence. The clone generating best fluorescence was chosen and designated pTOB.033.

### Integration and confirmation of plasmids pTOB.041 and pTOB.033 in HT-1080

Cells were engineered via electroporation (Lonza Nucleofector 2b), using Hanks Balanced Salt Solution (Sigma) as the electroporation buffer and the gRNA (Sigma) and Cas9 (Sigma) delivered as a Ribonucleoprotein (RNP). Cas9 solution (2 μl of 60 μM) and 5 μl of guide RNA hprt_sgRNA_2 (33 μM) were incubated together at room temperature for 10 minutes to form the RNP targeting *HPRT* intron 5. RNP (6 ul) was then added to an electroporation cuvette (Sigma) along with 2 μg of Donor Template plasmid and 1 × 10^6^ cells in prewarmed Hanks Balanced Salt Solution were then added to the cuvette and the cells electroporated using the L-005 program. Prewarmed Opti-MEM (300 ul; Gibco) was added to the cuvette then all of the cells were transferred to a six-well plate (Sarstedt) containing 3 ml of prewarmed complete media. Once the cells had recovered from electroporation, they were expanded up and then enriched for TagRFP658 by FACS. These enriched cells were then expanded. *HPRT* intron 2 was targeted with gRNA hprt_sgRNA_1 (33 μM) using the same protocol. Six μl of RNP was then added to an electroporation cuvette (Sigma) along with 2 μg of Donor Template plasmid and 1 × 10^6^ cells in prewarmed Hanks Balanced Salt Solution were then added to the cuvette and the cells electroporated using the L-005 program. 300 μl of prewarmed Opti-MEM (Gibco) was added to the cuvette then all of the cells were transferred to a 6 well plate (Sarstedt) containing 3 ml of prewarmed complete media. Once the cells had recovered from electroporation, they were expanded up and then enriched by FACS for TagRFP658 + and GFP+. Integration of the plasmids containing GFP or TagRFP658 constructs was confirmed by genomic DNA sequencing (Azenta) and alignment of the sequence reads spanning the plasmids with their DNA sequences (**Supplemental Fig. 2**).

### Fluorescence activated cell sorting (FACS) and flow cytometry

Prior to flow analysis or sorting, cells were detached, resuspended in a single cell suspension in FACS Buffer (PBS Sigma, 1% FBS Sigma, 0.5M EDTA Sigma, 1M HEPES Sigma) at a concentration of 1 × 10^6^ cells/ml. DAPI 1mg/ml (Sigma) was added as a live dead stain. Prior to sorting, cells were ran through a 40 μm filter (Fisher). A BD FACS Canto II was used for flow analysis with the DAPI excited by the 405 nm laser and measured in the 450/50 channel. GFP was excited by the 488 nm laser and measured in the 530/30 nm. TagRFP658 was excited by the 633 nm laser and measured in the 660/20 channel. A BD ARIA II was used for FACS.

### siRNA and small molecule inhibitor studies

siRNA duplexes were purchased from Sigma and resuspended in sterile nuclease free water to a final concentration of 100 μM (100 pmols/ul) and transfected using standard reverse transfection. For transfection of 1 × 10^6^ cells in a 10 cm cell culture dish, 20 μl of Lipofectamine® 2000 (Merck) was combined with 1 ml of Opti-MEM™ reduced serum medium (Merck). Separately, 200 pmols of siRNA duplexes (SiLuc 5’-CGUACGCGGAAUACUUCGA-3’ or siMSH3 5’-UCGAGUCGAAAGGAUGGAUAA-3’) were combined with 1 ml of Opti-MEM™ reduced serum medium. Following 5 minutes of incubation, the solutions containing Lipofectamine® 2000 and siRNA duplexes were gently combined and incubated for a further 15 minutes. Lastly 1 × 10^6^ cells in 9 ml of cell culture media were combined with the transfection solution and incubated for at least 48 hours. For 2-week long expansion assays, siRNA mediated knockdown was maintained by reverse transfection every 5 days.

### Western blotting

Cells were harvested for protein extraction using trypsinisation, washed using 1 X PBS and frozen as a dry pellet. Whole cell extracts were prepared using lysis in RIPA buffer followed by centrifugation. Protein concentrations of the resulting supernatant solutions were determined by the DC assay (BioRad) using bovine serum albumin as a standard. Extracts containing 60 ug total protein were separated on an 8% SDS-PAGE gel, transferred to nitrocellulose membranes, and probed with primary antibodies Msh3 primary antibody (BD Biosciences, 611390, mouse, 1/1000 dilution), Msh2 primary antibody (Calbiochem, NA26, mouse, 1/5000 dilution) or Msh6 primary antibody (BD Biosciences, 610919, Mouse, 1/1000 dilution). Incubation with Actin primary antibody (Sigma, A2066, Rabbit, 1/1000) and subsequent detection was performed simultaneously as a loading control. After overnight incubation and subsequent washing, membranes were washed and incubated with infrared-labelled secondary antibodies from Li-Cor Biosciences, IRDye^®^ 800CW goat anti-mouse IgG (P/N 925-32210) or IRDye^®^ 800CW goat anti-rabbit IgG (P/N 925-3211). Immunoreactive bands were visualised and quantified using Odyssey Infrared Imaging systems and software (Li-Cor Biosciences).

### Physical analysis of CAG repeat tract lengths

The DNA was isolated from these cells using a DNeasy blood and tissue kit (Qiagen). The DNA samples were quantified and diluted to a concentration of 50 ng/μl. The CAG repeat tracts were PCR amplified using the following primers: Forward Primer: 5’-TCAGCCTGGCCGAAAGAAAG-3’; Reverse Primer: 5’-GGTACCCGGGGATCCTCTAG-3’. The PCR products were size selected to remove primer dimers by using 0.6x and 0.8x AmpureXP bead purification. The purified samples were sent to the Shared Research Facility at the University of Glasgow for Illumina MiSeq sequencing. Fastq files obtained from the MiSeq sequencing were analysed using an alignment-based method on Galaxy. A reference file containing forward primers, reverse primers, flanking regions, and different CAG repeats was created. The sequences were aligned to this reference using the ‘BWA-MEM’ tool, with penalties for mismatches and gaps set according to Ciosi et al. [28]. Processed data were visualized as bar diagrams.

Capillary electrophoresis was performed by Cellmark Forensics on DNA extracted from the cells using DNAeasy blood and tissue kit (Qiagen). DNA was diluted to 10ng/μL and a PCR reaction mix was made using the following reagents: 10μL AmpliTaq™ Gold 360 Master Mix (2x) (4398881), 1μL forward primer (10μM, 5’-TCAGCCTGGCCGAAAGAAAG-3’), 1μL reverse primer (10μM, 5’-GGTACCCGGGGATCCTCTAG-3’), 1μL GC enhancer (supplied with AmpliTaq master max), 5μL water, 2μL gDNA (10ng/μL). 18μL of PCR reaction mix and 2μL of sample DNA extract were mixed and PCR was run according to the following conditions: 2 minutes at 95°C, followed by 28 cycles of 45s at 96°C, 45s at 59°C, 3 minutes at 72°C, followed by 1 minute at 72°C. Electrophoresis reaction mix was prepared with HiDi formamide (020-4001) and GeneScan™ 1200-LIZ Size Standard (4379950) in a ratio of 35:1. 1uL of PCR product and 9uL of reaction mix were mixed. DNA was denatured for 3 minutes at 95°C followed by 3 minutes at 4°C. Capillary electrophoresis was run in the 3500xL Genetic Analyser with a 36cm capillary array according to the ABI PRISM User Guide instruction suing GM IDXv1.5 software. The injection conditions were as follows: injection time 24s, injection voltage 1.2kV, run voltage 15kV, run temperature 60°C, and run time 2000s. All samples were analysed in duplicate on separate PCR runs, and the data was exported to a csv files and processed in JMP and R.

## Results

*TRX concept and creation of cell lines*. The purpose of the TRX assay is to measure CAG•CTG repeat expansions using a fluorescent readout to improve speed, sensitivity, and quantification. The assay is based on a human tissue culture line, HT-1080, a fibroblast line derived from a cancer patient. HT-1080 cells were chosen because they were shown to support CAG•CTG repeat instability in culture [29–31]. In addition, HT-1080 cells are near-diploid, they are easy to grow under biosafety level 1 conditions, they are amenable to experimentation, and they are suitable for flow cytometry analysis. The cells were obtained from an open repository (Leibniz Institute DSMZ). Simple tandem repeat analysis was used to confirm them as HT-1080 (**Supplemental Fig. 1A**).

The conceptual basis for the TRX assay is shown in **Fig. 1A**. HT-1080 cells were modified with two fluorescent markers. The green marker, d2GFP (hereafter, GFP), is expressed under the control of the Tet operator-repressor system [27] (**Fig. 1BC**). The Tet repressor protein is fused to the human KRAB corepressor to improve efficiency of silencing in human cells. If TetR-KRAB is expressed, it binds to the Tet operator and represses expression of GFP. However, the coding sequence of TetR contains an artificial intron, so correct splicing of the intron is required to express TetR-KRAB. The intron includes a CAG repeat tract. It is known that splicing efficiency of this intron is dependent on the length of the CAG repeat, with shorter tracts leading to more efficient splicing [23]. The starting repeat length allows splicing and adequate expression of TetR-KRAB to silence GFP (**Fig. 1B**). However, any cell in the population that undergoes a CAG repeat expansion is predicted to splice the intron less efficiently, leading to less repressor protein and relief of GFP silencing (**Fig. 1C**). The red marker, TagRFP658, is constitutively expressed (**Fig. 1D**) and serves as a normalization factor for fluorescence analysis. The net result of these events is that a CAG repeat expansion is predicted to convert the fluorescent profile of the cells from GFP – RFP + to GFP + RFP +.

Two TRX cell lines were established by integrating the red and green reporter plasmids into nearby sites within the *HPRT* gene on the X-chromosome. Since HT-1080 cells were derived from a male, there is only one X chromosome and therefore the two reporters are each in single copy. Genomic DNA sequence analysis confirmed single integrations at the desired loci (**Supplemental Fig. 2**). Sequence analysis also proved that the modified TRX cells were HT-1080, based on characteristic heterozygotic polymorphisms at *NRAS* and *IDH1* (**Supplemental Fig. 1B**). The two TRX cell lines were derived independently with the difference being the length and purity of the CAG repeat tract. The first line contains 63 repeats and is referred to here as (CAG)_63_. Sequencing analysis indicated that an aberrant interruption was present in repeat 15, such that the sequence is (CAG)_14_CTG(CAG)_48_. The interruption is presumably a cloning artifact. The second cell line contains 59 pure CAG repeats and is referred to as (CAG)_59_.

### Flow cytometry results from the TRX assay

The (CAG)_63_ cells were tested in the presence or absence of doxycycline (Dox) to exogenously control activity of the TetR-KRAB protein. When cells were cultured in the presence of Dox to inhibit TetR-KRAB function, the flow cytometry profile indicated a large majority of cells were GFP + RFP + (**Fig. 2A**, left panel). Double-positive cells (**Fig. 2A**, red box) were captured by FACS and the cells were then cultured in the absence of Dox to reestablish silencing of GFP. Flow cytometry analysis showed that almost all cells were GFP – RFP + (**Fig. 2A**, right panel). FACS sorting allowed establishment of a starting cell culture with the desired fluorescence profile. This double-sorting approach also minimizes false positive readings in subsequent steps of the assay. (CAG)_59_ cells respond similarly to this double-sorting procedure, and the approach was used routinely for all subsequent experiments.

**Figure 2.**
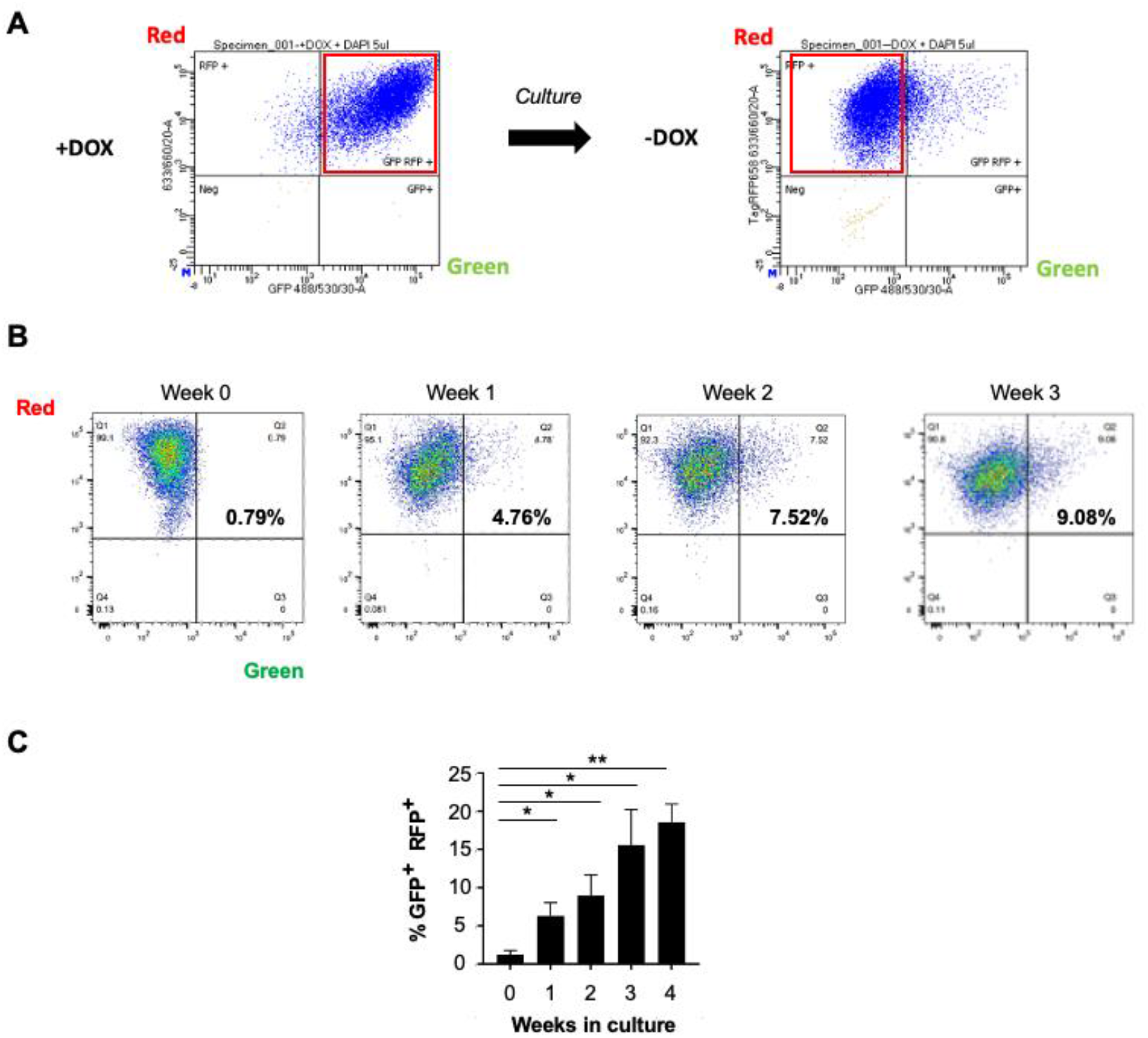
TRX assay results with (CAG)_63_ cells. **A**, FACS prior to onset of TRX assay. Cells were first grown in the presence of doxycycline to exogenously inactivate the TetR-KRAB repressor and allow GFP expression. Sorted cells (red box) were subsequently cultured without doxycycline to restore silencing of GFP and sorted for low green fluorescence (red box). The cells from the second sort were used for the TRX assay. **B**, Sample of flow cytometry data. Sorted cells from panel A were cultured for the indicated times. Flow cytometry analysis provided the percentage of GFP^+^ RFP^+^ cells indicated in each panel. **C**, Composite data from three biological replicates of three technical replicates each, including the data from **B**. Error bars denote ±SEM. *, *P*<0.05; **, *P*<0.001 by two-tailed Students t-test.

Sorted (CAG)_63_ cells were tested for the appearance of GFP + RFP + cells during culture. A sample data set is shown in **Fig. 2B**. The percentage of GFP + RFP + cells in the population increased steadily over several weeks in culture, from an initial reading of 0.79% to 9.08% after three weeks. Composite data for three independent biological replicates, each comprising three technical replicates, are shown in **Fig. 2C**. GFP + RFP + cells appeared at approximately 4-5% per week. Statistical analysis showed significance for each week 1-4 compared to week 0 (**Supplemental Table 1**). Based on this analysis, two weeks was selected as a standard assay length because it allows statistical confidence in a short timeframe.

### Assay validation

Validation of the TRX assay examined several key aspects. First, the independently derived (CAG)_59_ cell line was assayed (**Fig. 3A**). GFP + RFP + cells appeared at about 5% per week, like the rate seen for (CAG)_63_ cells. Since the repeat tract is pure in the (CAG)_59_ cells but interrupted in the (CAG)_63_ cells, the data support the idea that the interruption does not significantly change the rate of appearance of green cells in this assay. The second validation tested if the TRX signal depends on MSH3 protein. The prediction is that interfering with MSH3 should reduce expansions and therefore reduce the rate of appearance of TRX signal. MSH3 dependence was tested in two ways. Ablation of MSH3 abundance by siRNA was achieved that lasted for 15 days, the full length of the TRX assay. The average MSH3 knockdown efficiency was 82%, while levels of MSH6 and MSH2 were unchanged (**Fig. 3B**). TRX function was assayed and shown to be reduced by 51% compared to control level (**Fig. 3C**). This reduction is consistent with the interpretation that at least one-half of the TRX signal is MSH3-dependent, with the remaining signal presumably arising either from residual MSH3 protein or from non-specific events.

**Figure 3.**
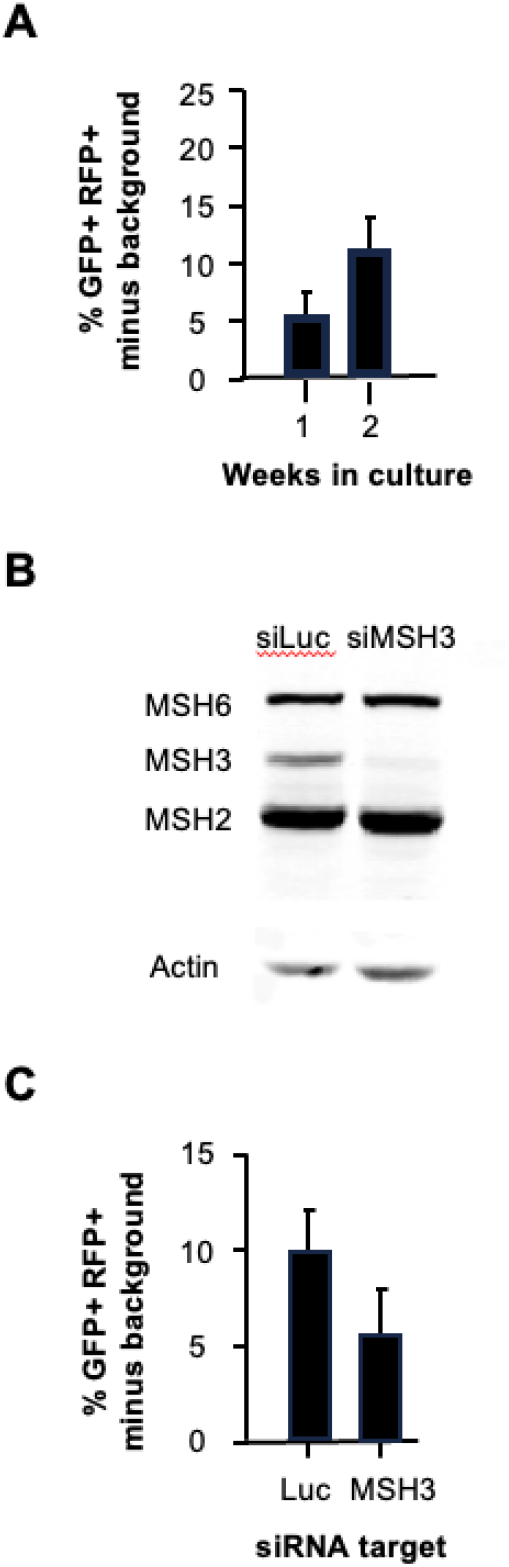
TRX assay validation. **A**, TRX assay results for (CAG)_59_ cells. Results are from three biological replicates of three technical replicates each. **B**, Western blot following siRNA knockdown of either luciferase or MSH3. Cells were transfected with siRNA at days 0, 5 and 10. Protein lysates were prepared at day 15. Total protein (60 ug) was analyzed in each lane. Three iterations yielded an average MSH3 knockdown efficiency of 82% ± 4%. **C**, TRX assay results at day 15 of the siRNA knockdown for three biological replicates. Error bars denote ±SEM. The average reduction in TRX activity upon MSH3 ablation was 51% ± 14%.

### Physical analysis of the CAG repeat tract lengths

The CAG repeat tracts were examined in TRX cell populations by PCR followed by ultradeep MiSeq sequencing (**Fig. 4AB**). The samples shown in the figure correspond to the three biological replicate cell cultures from **Fig. 2C**. Sequencing analysis showed that all three cultures started with a modal repeat length of 63. There were some smaller peaks with sizes from about 60-64 repeats, likely due to PCR polymerase slippage. The validity of the approach was confirmed as shown in the table view (**Fig. 4B**), where the CTG interruption at triplet 15 is clearly visible. In all, the starting cultures gave length distributions that are expected from this technique. The spectra after four weeks of culture did not show obvious CAG•CTG repeat expansions. Rather, in many (but not all) of the cultures, a second population of peak size centred around 29 repeats was observed. The source of the 29 repeat signal was not obvious. We tested many different cultures, including those which were sorted for high green fluorescence and which should, in principle, be enriched for expansions according to the scheme in **Fig. 1**. In none of the cultures did we see good evidence for expanded alleles at the frequency that would generate the corresponding levels of green fluorescence. Either the spectra of the finishing cultures looked much like those from the starting culture, or there was evidence of shorter alleles like in **Fig. 4A**.

**Figure 4.**
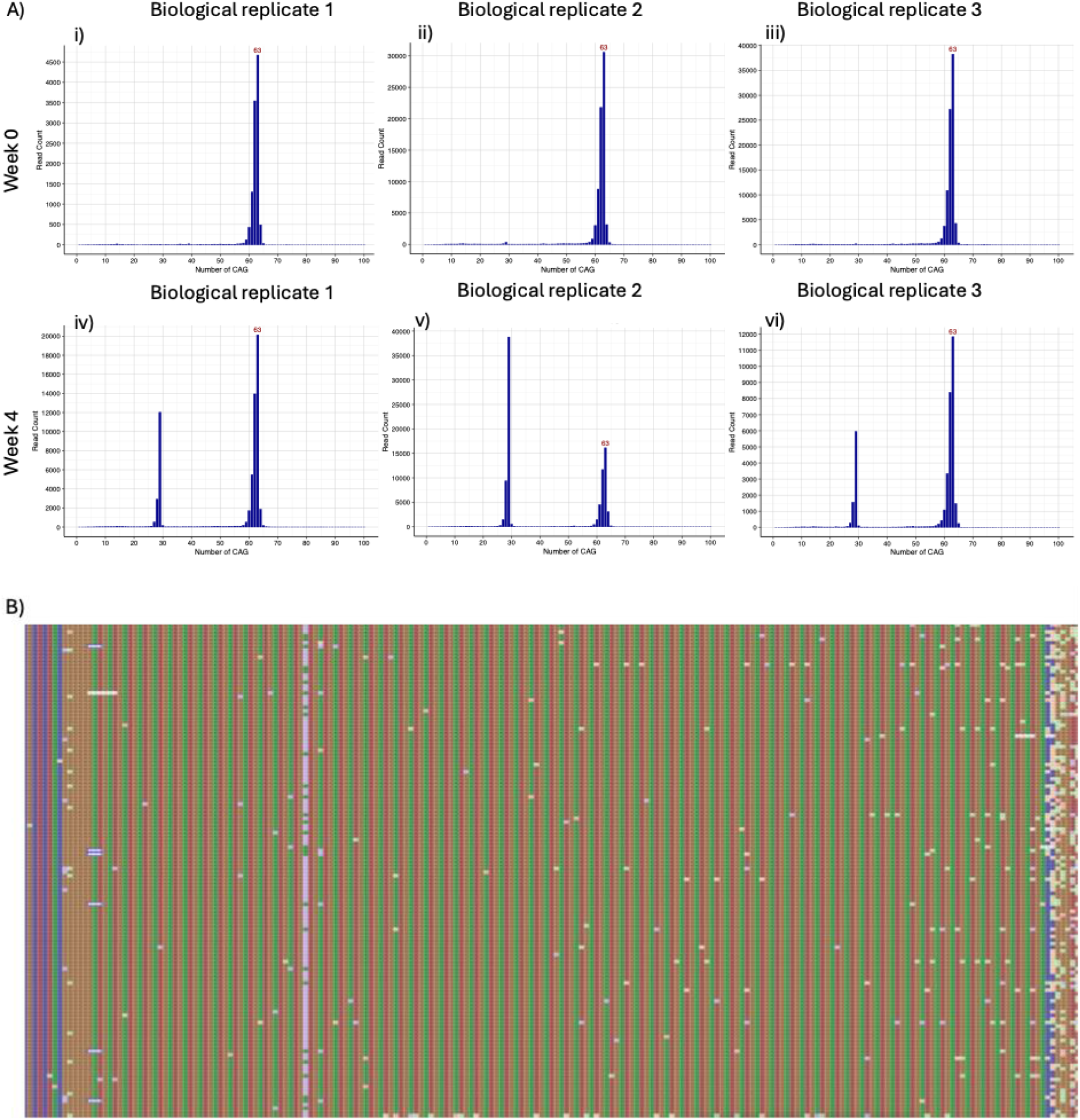
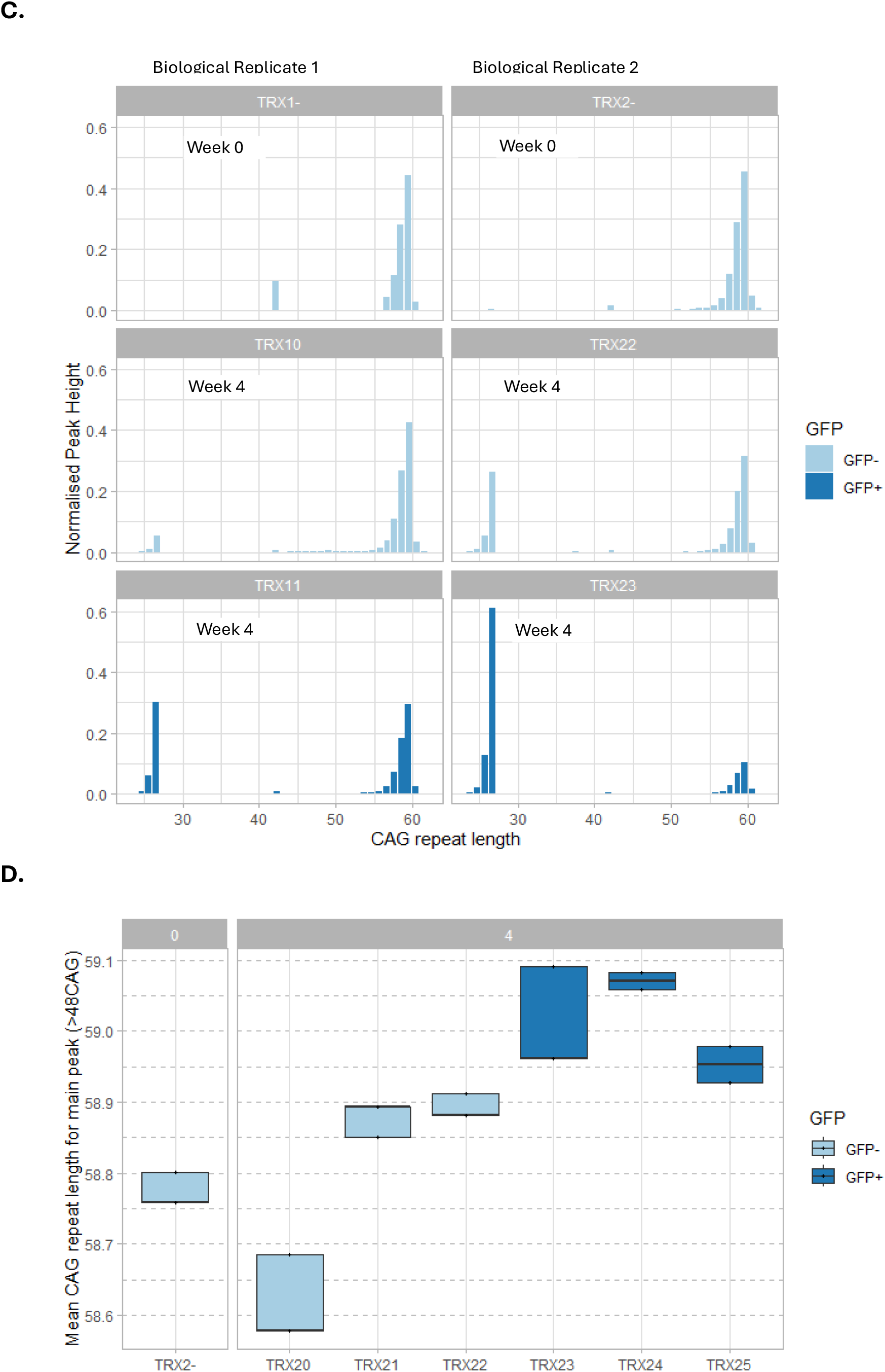
Analysis of CAG•CTG tract lengths by MiSeq and capillary electrophoresis. **A**, MiSeq bar graphs of the CAG•CTG sizes plotted against the number of reads. Three biological replicates at week 0 and week 4 (high GFP fluorescence). **B**, MiSeq tablet view of the (CAG)_63_ repeat tract with a CTG interruption at repeat 15. **C**, Analysis of CAG tract lengths by capillary electrophoresis. CAG tract lengths shown for two biological replicates in cells sorted for GFP+ or GFP- and at two time points. **D**, Mean CAG repeat length for the major peak in biological replicate 2 samples. Major peak data was isolated (CAG>48), and the mean repeat length was calculated using the normalised peak height.

CAG repeat tract lengths measured using capillary electrophoresis showed similar profiles to those produced using MiSeq sequencing. **Fig. 4C** shows some example traces from week 0 and week 4 cells sorted for GFP- or GFP+. Week 0 samples showed a modal repeat length of 60, which differs from that determined by MiSeq, probably due to the reduced resolution of capillary electrophoresis. We also observed peaks in most of the samples around 27 repeats, and in some samples a small intermediate peak around 42 repeats. We did not observe any significant expansions in any of the samples, however the mean CAG repeat length for the major peak (using 48 repeats as a lower limit) did appear slightly greater higher for GFP+ samples (**Fig. 4D**), although this difference was not significant.

## Discussion

The TRX assay provides the framework for more rapid and more sensitive detection of CAG•CTG repeat expansions, compared to current assays. The low green fluorescence background makes it feasible to readily detect increases in signal, and the inclusion of the red fluorescent marker increases the robustness of the assay. Using standard FACS and flow cytometry, the TRX assay yielded the predicted increase in green fluorescence rapidly (one-two weeks) and with good sensitivity of <2%. These features are significant improvements over current assays for expansions, which typically require ∼six weeks of culture and are not as sensitive. The availability of two independently derived TRX lines, containing 63 CAG repeats with one interruption or 59 pure CAG repeats, offers useful validation options. Additional validation took advantage of the known dependence of repeat expansions on the MSH3 subunit of the DNA repair protein MutSβ. Knockdown of MSH3 reduced the TRX signal, with residual signal likely due to incomplete knockdown/inhibition or to unrelated genetic events that produce background fluorescence.

The CAG•CTG tract lengths in the TRX assay fall into a size range that is particularly relevant for somatic expansions in Huntington’s disease. It was recently demonstrated [7] that somatic expansions appear to be a required step in neuronal cell death in HD, and that the window of somatic expansions prior to death is quite large (up to ∼150 repeats). Furthermore, somatic expansions appear to fall into two categories, a ‘slowly ticking clock’ that increases the inherited repeat length of 40-50 repeats up to about 80 repeats, followed by a more rapid expansion phase from about 80 to >150 repeats. Our assay uses allele lengths of 59 and 63 repeats, within the crucial ‘slowly ticking clock’ timeframe where therapeutic intervention might be particularly fruitful in delaying or blocking the onset of HD.

Another virtue of the TRX assay is technical simplicity. With the TRX cells in hand, the assay requires only standard tissue culturing, FACS and flow cytometry. The fluorescent readouts are well within common detection parameters available on many cytometers, and standard software allows ready quantitative readouts. The HT-1080 cells are amenable to standard manipulations, such as siRNA or small molecules, provided that the treatment does not alter cell growth or viability in any major way. Gene editing techniques such as CRISPR-Cas9 have also been performed successfully in HT-1080 cells herein and, for example, [32,33], which should allow standard gene knockout or other genetic manipulations in the TRX system.

The cell sorting protocol shown in **Fig. 2A** was very useful in minimizing false positive and false negatives cells for all experiments. The results in **Fig. 2BC** were obtained by starting the assay immediately after sorting. This resulted in a low background level of 0.79%. We also tried an alternative protocol where cells were sorted, then frozen in aliquots. Subsequent thawing and recovery growth occurred without further sorting. The thawing and cell growth recovery took approximately five to seven days, during which time some cells developed higher green fluorescence which put them into the GFP+ range. In other words, there was substantial background of 4-10% prior to the onset of the experiment. While this background was measured and accounted for in the TRX assay results shown in **Fig. 3ACD**, the higher levels led to more scatter in the data and, correspondingly, higher *P* values (**Supplemental Table 1**). As an improvement going forward, we recommend that all TRX assays initiate immediately following the sorting protocol shown in **Fig. 2A**. This improvement should reduce scatter in the data and improve statistical confidence.

Surprisingly, physical analysis of the CAG repeat tracts by ultradeep MiSeq or by genescan approaches showed no significant level of expansions, even in cell populations with enhanced green fluorescent signal (**Fig. 4**). We do not fully understand why expansions are infrequent in these cells, and this shortcoming must be overcome with additional research to maximize the utility of the TRX assay. Overall, the development of the TRX assay provides a useful platform for further exploration to better link the TRX fluorescent signal with verifiable triplet repeat expansions.

## Acknowledgement

This project received funding from the European Union’s Horizon 2020 research and innovation programme under the EJP RD COFUND-EJP N°825575. Additional funding was from LoQus23 Therapeutics Ltd. VD is further supported by the UK Dementia Research Institute (award number UK DRI-3006) through UK DRI Ltd, principally funded by the Medical Research Council. The authors gratefully acknowledge Dr Waheba El-Sayed of UK Dementia Research Institute at Cardiff University for providing the (CAG)_123_ PCR template and Dr Shirley Hanley of the University of Galway Flow Cytometry Core Facility for expert guidance.

## Notes

### Competing Interest Statement

Paulina Marzec, Madeleine Richer and Caroline Benn are employees of LoQus23 Therapeutics Ltd, Cambridge, UK.
Robert Lahue is a paid consultant of LoQus23 Therapeutics Ltd, Cambridge, UK.

