## Supplemental information for "The TRX assay for triplet repeat expansions"

**Supplemental Table 1**

n and *P* values

All *P* values from two-tailed Student's t-test

| Figure | Entry | Comparator | n | *P* |
|---|---|---|---|---|
| 2C | GFP+ RFP+ @1 wk | GFP+ RFP+ @ 0 wk | 3* | 0.035 |
| 2C | GFP+ RFP+ @2 wk | GFP+ RFP+ @ 0 wk | 3* | 0.032 |
| 2C | GFP+ RFP+ @3 wk | GFP+ RFP+ @ 0 wk | 3* | 0.043 |
| 2C | GFP+ RFP+ @4 wk | GFP+ RFP+ @ 0 wk | 3* | 0.0003 |

*Three biological replicates of three technical replicates each

| | | | | |
|---|---|---|---|---|
| 3A | GFP+ RFP+ @1 wk | GFP+ RFP+ @ 0 wk | 3* | 0.18 |
| 3A | GFP+ RFP+ @2 wk | GFP+ RFP+ @ 0 wk | 3* | 0.70 |

*Three biological replicates of three technical replicates each

| | | | | |
|---|---|---|---|---|
| 3A | GFP+ RFP+ siMSH3 | GFP+ RFP+ siLuc | 3* | 0.30 |

*Three biological replicates

**Supplemental Table 2**

PCR primers and guide RNAs

PCR primers

| | |
|---|---|
| oTOB.029 | atcatgaatcttcttgtggcagtg |
| oTOB.032 | ggctgttagtgataagtgatcagga |
| oTOB.105 | gcatcaagtcgctaaagaagGTAGGTAATTTGAGAAAAGC |
| oTOB.106 | agtagtaggtgtttccctttCTGCATTCAGAAGAATAAGAG |
| oTOB.109.02 | GAGATAGTCTGCAAATTCAGTGATGC |
| oTOB.109.03 | AGCATGTGTTTATCTAGCGCTG |
| oTOB.144 | catcctaaaggtaaggcTAGTTATTAATAGTAATCAATTACGGG |
| oTOB.145 | tttctctccctgggcACGCCTTAAGATACATTG |
| oTOB.146 | ggaactccatatatgggctatgaac |
| oTOB.148 | gactacagactggaaagaatcgagg |
| 040.05 | caaagaccccaacgagaagc |
| 040.06 | catggtaatagcgatgactaatacgtag |

Guide RNAs

| | | |
|---|---|---|
| gRNA7 | GCUUCCCUUUACACAACGUU | |
| gRNA8 | AAAUAUCAACAAAACAUCAG | |
| hprt1_sgRNA_1 | GCAUUUCUCAGUCCUAAACA | targeting to *HPRT* site 1 |
| hprt1_sgRNA_2 | ACCAUCCUAAAGGUAAGCCA | targeting to *HPRT* site 2 |

### Supplemental figures

**A**

**Result:**

| **Client Sample Name** | HT-1080 WCB |
|---|---|
| **Sample Code** | CL00009584 |
| D8S1179 | 13,14 |
| D21S11 | 28,30 |
| D7S820 | 9,10 |
| CSF1PO | 12,12 |
| D3S1358 | 16,16 |
| TH01 | 6,6 |
| D13S317 | 12,14 |
| D16S539 | 9,12 |
| D2S1338 | 25,26 |
| D19S433 | 13.2,15 |
| vWA | 14,19 |
| TPOX | 8,8 |
| D18S51 | 12,18 |
| AMEL | X,Y |
| D5S818 | 11,13 |
| FGA | 22,25 |
| **Database Name** | HT-1080 |

**B**

**Expected* and Observed****

| | | Gene sequence | Protein sequence |
|---|---|---|---|
| Chr1 | *NRAS* | heterozygous c.181C>A | Q61K |
| Chr2 | *IDH1* | heterozygous c.394C>T | R312C |

**Supplemental Figure 1.** Genetic validation of HT-1080 cells

**A**, Simple tandem repeat analysis of the HT-1080 cells prior to addition of reporter plasmids. Data provided by a commercial vendor (https://eurofinsgenomics.eu/).

**B**, DNA sequence analysis of HT-1080 specific variants in both (CAG)63 and (CAG)59 cell lines.

*Expected, ATCC [https://www.atcc.org/products/ccl-121].

**Observed, from genomic DNA sequencing [https://www.genewiz.com]

**Supplemental Figure 2.** Confirmation of plasmid integrations into *HPRT*

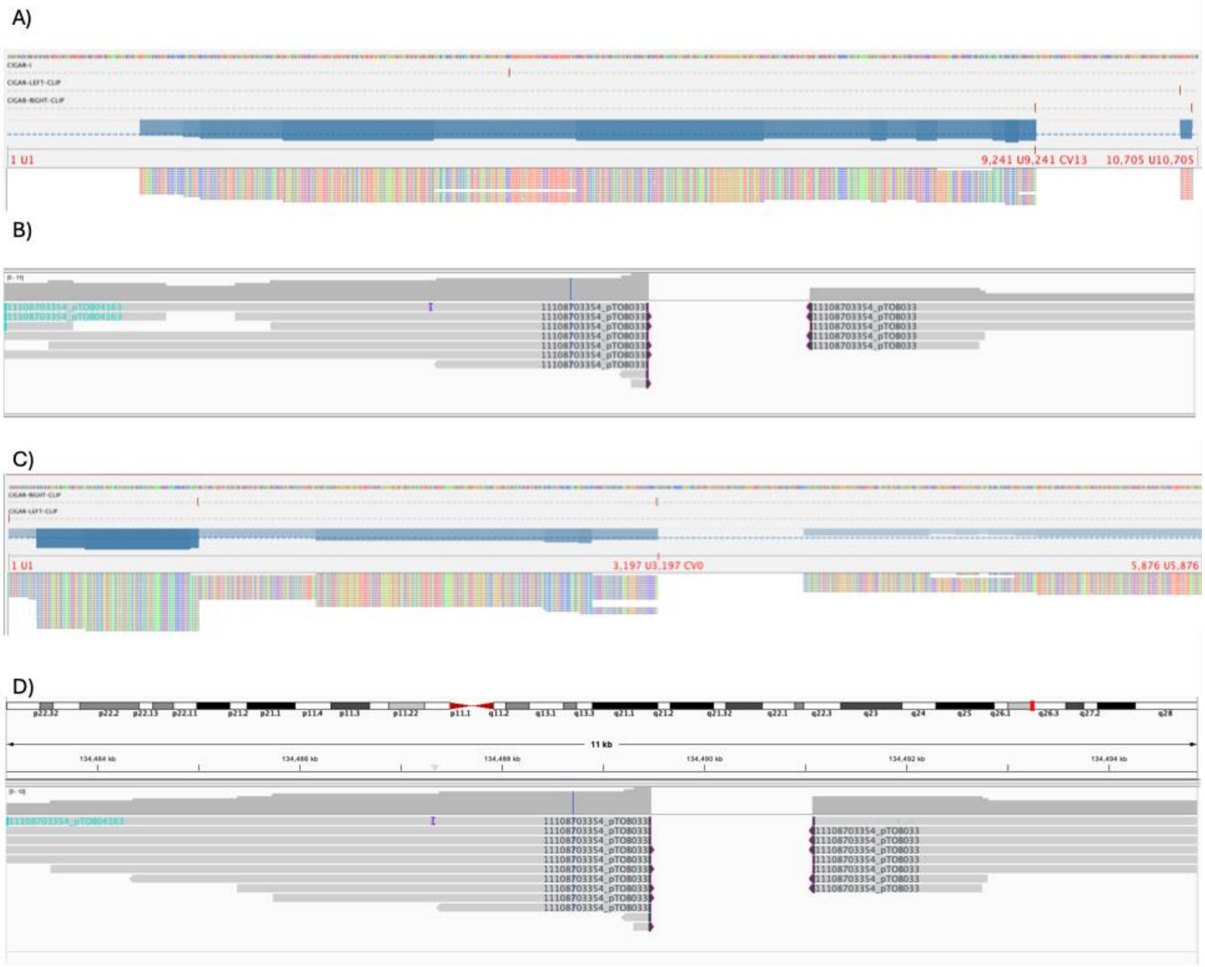


**Supplemental Figure 2.** Confirmation of plasmid integrations into *HPRT*

**A**, Tablet view of the sequenced plasmid aligned to the reference plasmid sequence (pTOB04163).

**B**, Genomic DNA analysis showing the reads aligning to the *HPRT* gene showing deleted chrX:134,472,947 to 134,474,548, replaced by pTOB04163.

**C**, Tablet view of the sequenced plasmid aligned to the reference plasmid sequence (pTOB033).

**D**, Genomic DNA analysis showing the reads aligning to the *HPRT* gene showing deleted chrX: 134,489,477 to 134,491,077, replaced by pTOB033.
